# Detecting recent selective sweeps while controlling for mutation rate and background selection

**DOI:** 10.1101/018697

**Authors:** Christian D. Huber, Michael DeGiorgio, Ines Hellmann, Rasmus Nielsen

## Abstract

A composite likelihood ratio test implemented in the program SweepFinder is a commonly used method for scanning a genome for recent selective sweeps. SweepFinder uses information on the spatial pattern of the site frequency spectrum (SFS) around the selected locus. To avoid confounding effects of background selection and variation in the mutation process along the genome, the method is typically applied only to sites that are variable within species. However, the power to detect and localize selective sweeps can be greatly improved if invariable sites are also included in the analysis. In the spirit of a Hudson-Kreitman-Aguadé test, we suggest to add fixed differences relative to an outgroup to account for variation in mutation rate, thereby facilitating more robust and powerful analyses. We also develop a method for including background selection modeled as a local reduction in the effective population size. Using simulations we show that these advances lead to a gain in power while maintaining robustness to mutation rate variation. Furthermore, the new method also provides more precise localization of the causative mutation than methods using the spatial pattern of segregating sites alone.

## Introduction

Rapid advances in sequencing technology during the past few years have facilitated studies using genome-wide molecular data for detecting signatures of selective sweeps (Akey *et al.* 2002; Carlson *et al.* 2005; Wang *et al.* 2006; Voight *et al.* 2006; Kelley *et al.* 2006; Kimura *et al.* 2007; Williamson *et al.* 2007; Tang *et al.* 2007; Sabeti *et al.* 2007; Xia *et al.* 2009; Qanbari *et al.* 2012; Ramey *et al.* 2013; Long *et al.* 2013; Chávez-Galarza *et al.* 2013; Huber *et al.* 2014), and a large number of computational methods have been developed for this purpose (e.g., Fu & Li 1993; Kim & Stephan 2002; Sabeti *et al.* 2002, 2007; Kim & Nielsen 2004; Nielsen *et al.* 2005; Voight *et al.* 2006; Jensen *et al.* 2007; Boitard *et al.* 2009; Chen *et al.* 2010; Pavlidis *et al.* 2010, 2013; Li 2011). The various methods differ in the assumptions that they make about the selective sweep. For example, the extended haplotype test and its derivatives are powerful in cases where the beneficial mutation has not yet reached fixation in the population (Sabeti *et al.* 2002, 2007; Voight *et al.* 2006). Methods based on measures of population subdivision rest on the assumption that in geographically structured populations a selective sweep has a locally confined effect on genetic diversity, which increases population differentiation at the position of the sweep (Akey *et al.* 2002; Chen *et al.* 2010). More recently, statistics have been developed specifically for the detection of soft sweeps, i.e. a pattern caused by multiple haplotypes sweeping to high frequencies (Ferrer-Admetlla *et al.* 2014; Garud *et al.* 2015).

In the present study, we are solely concerned with the model of a classical hard selective sweep in a single population, and we assume that the beneficial mutation has reached fixation not too long ago. The methods usually applied in this scenario aim to detect deviations in the shape of the site frequency spectrum (SFS), which can be quantified with simple summary statistics like Tajima’s D or Fay and Wu’s H. In addition, more powerful statistics have been developed that explicitly model the effect of a selective sweep on the SFS in a likelihood ratio framework (Kim & Stephan 2002; Nielsen *et al.* 2005). Kim and Stephan (2002) proposed a composite likelihood ratio statistic based on calculating the product of marginal likelihood functions for all sites on a chromosome under models with and without a selective sweep at a particular position, and under the assumption of a panmictic population of constant size. The resulting composite likelihood ratio is then computed for each position of interest to evaluate the evidence for a sweep at those positions. This method, therefore, does not only incorporate information regarding the SFS, but does so in a way that uses the spatial distribution of segregating alleles of different frequencies. The null distribution of the test statistic is approximated using simulations. An extension to this test was proposed by Nielsen *et al*. (2005). In this method, the overall genomic SFS is used as the neutral, or background, model instead of using the standard neutral model as the null. The distribution of the SFS under the alternative hypothesis of selection is derived by considering the way a selective sweep would modify the observed background distribution of allele frequencies. This leads to a computationally fast method, facilitating genome-wide analyses. Nielsen *et al.* (2005) also argued that the use of the overall genomic SFS to represent the neutral case leads to increased robustness, and showed that the method was robust to a two-epoch growth model and an isolation-migration model with population growth in both populations, with parameters estimated from human single nucleotide polymorphism (SNP) data (Marth *et al.* 2004). Since then, it has become clear that while this method may be more robust than some previous SFS-based approaches, it can produce a high proportion of false positives if there has been a strong recent bottleneck in population size, but a standard neutral model is used to calculate critical values (Jensen *et al.* 2005; Pavlidis *et al.* 2008).

If invariable sites are included in the analysis, then both the methods of Kim and Stephan (2002) and Nielsen et al. (2005) may be sensitive to assumptions regarding selective constraint and mutation rates. A region with strongly reduced levels of variation due to selective constraint or reduced mutation rate may be misinterpreted as a region that has experienced a recent selective sweep (Nielsen et al., 2005; Boitard et al., 2009; Pavlidis et al., 2010). For these reasons, Nielsen et al. (2005) proposed to use only polymorphic sites, an option that became incorporated as default in both Sweepfinder (Nielsen *et al.* 2005) and SweeD (Pavlidis *et al.* 2013).

Background selection can also lead to locally reduced levels of neutral variation (Charlesworth et al., 1993, 1995; Hudson and Kaplan, 1994, 1995; Nordborg et al., 1996; Charlesworth, 2012; Cutter and Payseur, 2013) and cannot be ignored for the study of neutral polymorphisms in many cases (Williford & Comeron 2010; Cutter & Payseur 2013; Messer & Petrov 2013). Cutter and Payseur (2013) argue that the inevitability and prevalence of deleterious mutations necessitates the incorporation of background selection in the null model when identifying positive selection. There is a well-developed mathematical framework for quantifying the strength of background selection given the genome-wide mutation rate, recombination rate, position of functional elements and distribution of fitness effects (Hudson & Kaplan 1995; Nordborg *et al.* 1996). As data sets and methods for estimating the effect of background selection for each position in the genome are becoming available (McVicker *et al.* 2009; Comeron 2014), the objective of developing methods for detecting positive selection that can take background selection into account is becoming tenable.

Here, we explore the potential for improving the composite likelihood ratio test of SweepFinder (Nielsen *et al.* 2005) by either including invariant sites that differ with respect to an outgroup (*i.e.*, fixed differences), or all invariant sites, in addition to polymorphic sites. When only including fixed differences, the method incorporates the information typically represented in a Hudson-Kreitman-Aguadé (HKA) test (Hudson *et al.* 1987), but adds the information from the spatial distribution of allele frequencies. We show that this approach is robust to variation in mutation rate across the genome, and develop an approach for incorporating estimates of the strength of background selection into the SweepFinder framework. Using the reduction in diversity relative to divergence as a necessary hallmark of a selective sweep in our model also helps to reduce false positives, *e.g.* in the case of a recent population bottleneck. Finally, we compare results of both the old and the new version of the likelihood ratio test applied to human genetic data.

## Materials and Methods

### Including invariant sites into the SweepFinder framework

Starting with *n* aligned DNA sequences, each of length *L*, we wish to determine whether a selective sweep has occurred at some defined position along the sequence. Based on results of Durrett and Schweinsberg (2004), Nielsen et al. (2005) derived an approximate formula for *p*_*k*_*, the probability of observing *k* derived alleles, *k* ∈{1,2,…,*n*- 1}, in a sample of size *n*, immediately after a selective sweep, for a site at a particular distance (*d*) from the selected mutation. For each *k*, *p*_*k*_* is a function of *d*, the background allele frequency distribution **p**=(*p*_1_,*p*_2_,…,*p*_*n*-1_), and the parameter α=*r* ln(2 *N*_*e*_)/*s*. Here, *r* is the per-base per-generation recombination rate, *s* is the selection coefficient, and *N*_e_ is the effective population size. The parameter *p*_*k*_ is the expected proportion of sites, not affected by the sweep, in which the derived allele has a frequency of *k*/*n* in the sample. The vector **p** is commonly estimated as the observed SFS from the whole genome, under the assumption that only a small and therefore negligible proportion of positions are affected by selection. The parameter α quantifies the relative influence of recombination and selection, with small values of α indicating strong sweeps.

The equations in Nielsen et al. (2005) allow for the incorporation of invariant sites that may or may not be fixed differences relative to an outgroup, using **p**=(*p*_0_, *p*_1,_…,*p*_*n*_) as the definition of **p**, and with the modification that the upper limit of the sum in equation (5) of Nielsen et al. (2005) is *n* and not *n*-1. The quantity *p*_*k*_* is a function of the probability of a lineage escaping a selective sweep, *P*_e_=1-exp(-α*d*), where *d* is the distance between the polymorphic site and the sweep location. Our new version of SweepFinder allows defining distances between sites as genetic distance. This is achieved by allowing *d* to be defined by a recombination map rather than by physical distance as in the previous version. As in Nielsen et al. (2005), we then define the composite likelihood ratio statistic CLR = 2[log(CL_sweep_) - log(CL_background_)], where CL_sweep_ is the composite likelihood maximized over alpha, and CL_background_ is the composite likelihood calculated under the assumption of α = ∞. This is a composite likelihood ratio, and not a full likelihood ratio, because sites in the genome are not independent, but correlated due to linkage disequilibrium. One thing to notice, about which there has existed some confusion in the literature, is that this approach is not window based but in theory incorporates information from all SNPs in the genome to inform the CLR calculated for a single point in the genome. However, for computational efficiency SweepFinder uses a cut-off for distances from the focal SNP to include in the calculation. As distances become large, the contribution to the likelihood ratio approaches zero. The value used for the cut-off in SweepFinder is αd = 12, corresponding to a probability of a lineage escaping a sweep of 0.999994. Furthermore, SweepFinder calculates probabilities on a grid of recombination distances and uses a smooth interpolation to approximate probabilities for a particular point.

The effect of including invariant sites on the SFS is illustrated in Figure 1. In a region close to the site of the selective sweep, variability is reduced because almost all the probability mass is concentrated on fixed alleles. Notice also, that as the mutation rate scales all categories proportionally, a change in the mutation rate will not change the SFS defined on {1, 2,…,*n*} (Figure 1a and 1b). This statement does not hold true when invariant sites that do not differ from the outgoup (Figure 1c) are incorporated.

**Figure 1.**
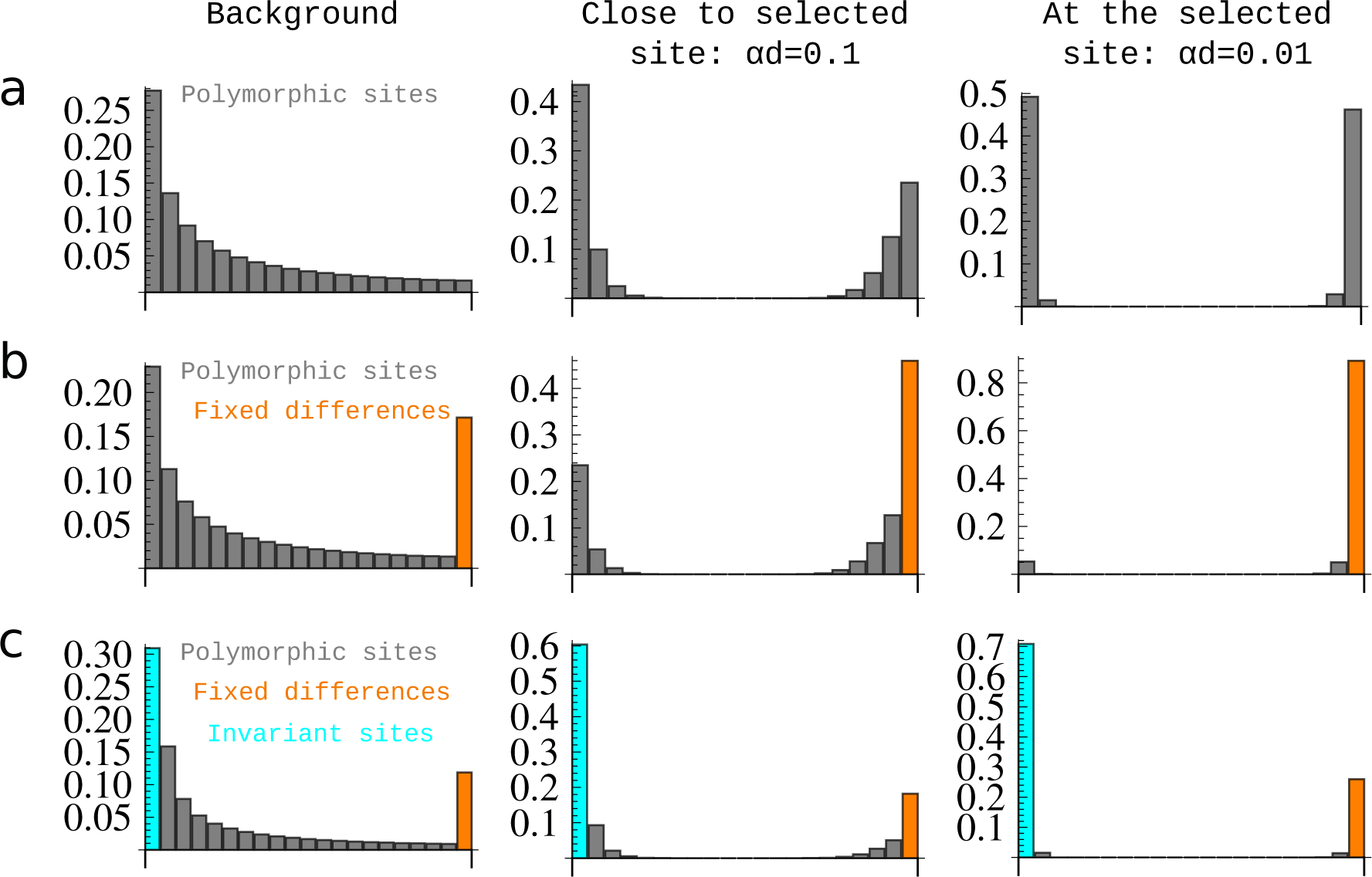
Effects of a selective sweep on the expected SFS. a) The expected SFS of a standard neutral background and of a neutral site linked to a selective sweep, assuming different distances between the neutral site and the sweep locus. b) The same expectations for a SFS that is extended to include the class of fixed differences (sites that are invariant in the sample, but different to an outgroup species). c) The same expectations for a SFS that is extended for the class of fixed differences and invariant sites that do not differ from the outgroup species. All expectations are calculated with the formulas in Nielsen et al. (2005).

### Accounting for background selection

A B-value (*B)*, is the factor by which the effective population size is expected to be reduced due to background selection, *i.e. N*_*e*_* = *N*_*e*_*B*, where *N*_*e*_ and *N*_*e*_* are the effective population sizes with and without background selection, respectively (Charlesworth 2012). We will assume that a reasonable estimate of the ‘B-value map’, the value of *B* for each site in the genome, is available (see for example McVicker et al., 2009, for humans). We note that this assumption limits the use of our method to organisms for which such estimates have been obtained. We also note that we only model the main effect of background selection: the well-known reduction in effective population size. However, background selection can also affect the distribution of allele frequencies (Charlesworth *et al.* 1993, 1995; Hudson & Kaplan 1994; Zeng & Charlesworth 2011; Lohmueller *et al.* 2011a; Nicolaisen & Desai 2013), an effect that is ignored here.

Based on the B-value map, the expected site frequency spectrum can be adjusted simply by multiplying all categories in the spectrum, except for the zero and the *n* (fixed differences) category, by *B*, *i.e.*, by setting 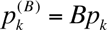 for *1* < *k* < *n-1*, as the expected diversity reduction is proportional to *B*. The *n* category can be adjusted as described in the next section. The zero category can be obtained by standardization, *i.e.* 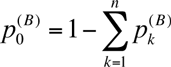. If the zero category is not included in the analysis, all included categories will have to be standardized to ensure that the frequencies sum to 1. The calculation of the CLR then proceeds as in Nielsen *et al.* (2005).

### Effect of background selection on number of fixed differences

We assume the availability of a sample of *n* chromosomes and a single chromosome from an outgroup species, which split from the ingroup species *g* generations ago. Fixed differences are defined as sites with an allele that is invariant within the ingroup sample, but different from the allele at the orthologous position of the outgroup chromosome (Figure 2). The expected number of fixed differences, *K*, in the sample is then *E*[*K*] = *μ*(2*T*_*anc*_ − *T*_*in*_), where *T*_*in*_ is the time to the most recent common ancestor in the ingroup sample, *T*_*anc*_ is the divergence time between ingroup and outgroup, and *μ* is the per-generation mutation rate. We further assume a standard neutral coalescent model with populations of constant sizes *N*_*e,in*_ and *N*_*e,anc*_ for the ingroup population, and ancestral population, respectively (Figure 2), and that the split time, *g,* is so large that we can assume Pr(*T*_*in*_ > *g*) ≈ 0. Then E[*T*_*in*_] = 4*N*_e,in_(1-1/*n*), where *n* is the sample size of ingroup sequences, and E[*T*_*anc*_] = g+2*N*_e,anc_. Then, under an infinite sites model

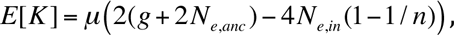

and the relative number of fixed differences with and without background selection is

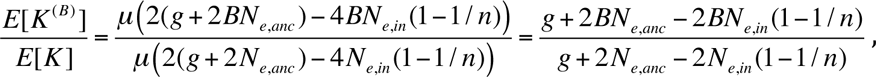

which reduces to

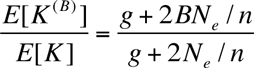

for *N*_e,anc_ =*N*_e,in_ = *N*_e_. In the limit of large split times (*g* → *∞*), E[K^(B)^]/E[K] ≈ 1, and the effect of background selection on fixed differences can generally be ignored if *g* >> *N*_*e*_/*n*. In our new version of SweepFinder, if the B-value map is included for sweep detection, estimates of *N*_*e,in*_, *N*_*e,anc*_, and *g* have to be provided to the software.

**Figure 2.**
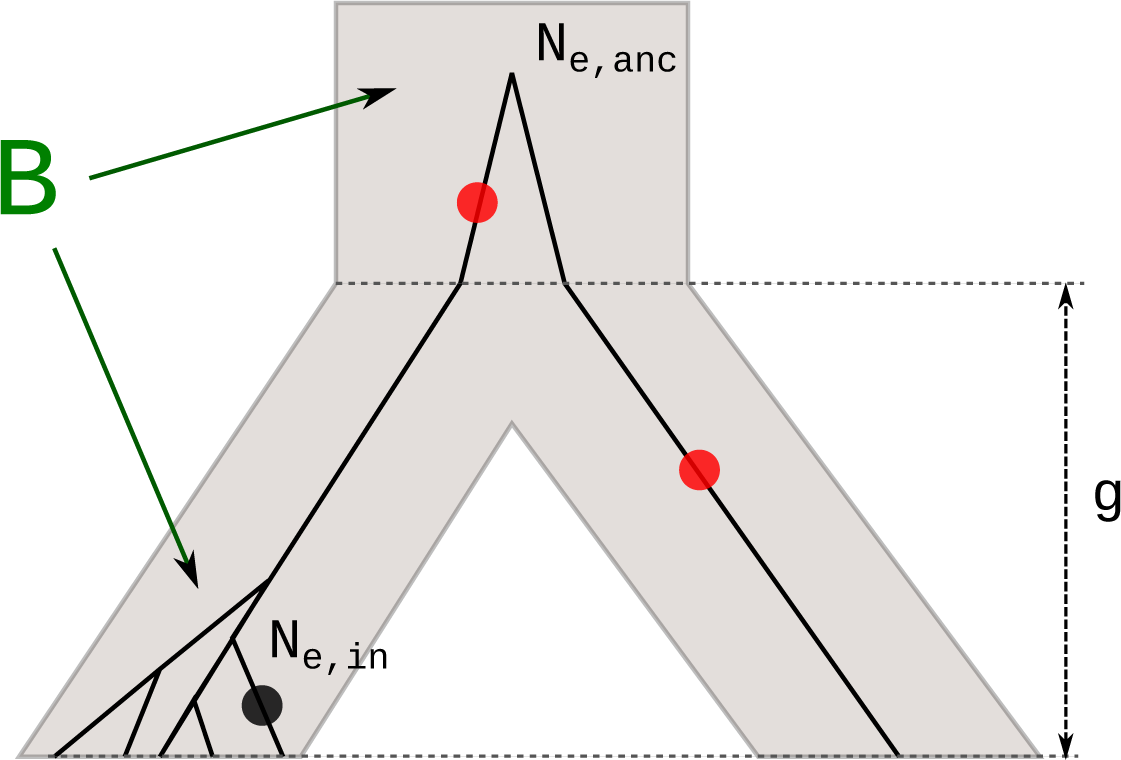
Definition of fixed differences and polymorphic sites. We define fixed differences (red) as sites that are not polymorphic within the ingroup and differ between ingroup and outgroup. Note that mutations on the lineage to the outgroup also count as a fixed difference. Here, the ingroup is sampled with 5 chromosomes, the outgroup with one chromosome. Background selection influences both the number of fixed differences and the number of polymorphisms

### Constant size and bottleneck simulations

Simulations were performed under the model described in Figure 2 assuming *L* = 100kb and *n* = 30, using *msms* (Ewing & Hermisson 2010).

We set the split time, *g*, between ingroup and outgroup to 20 coalescent time units (2*N*_*e*_ generations), resulting in a neutral divergence of 0.1. The scaled mutation rate θ = 4*N*_*e*_μ per site was set to 0.005 and the population scaled recombination rate per site, 4*N*_*e*_r, to 0.02. Those parameters where chosen to be comparable to the ones in (Nielsen *et al.* 2005). One chromosome was sampled from the outgroup species to classify invariant sites into sites that differ or do not differ to the outgroup. To analyze the effect of reduced mutation rate in a genomic region compared to the background, we varied the mutation rate between 0.1 and 0.9 times the mutation rate in other regions. Further, we simulated two demographic scenarios, a constant size and a bottleneck population. In simulations with selection, the selected mutation was introduced in the population at specified times (0.01, 0.02, 0.04, 0.06, 0.08, 0.16, 0.24,…, 1.2), at a frequency of 1/(2*N*_*e*_ with a population scaled selection coefficient of 2*N*_e_s = 200. We only kept simulations in which the mutation did not get lost (-SFC option in *msms*).

For the bottleneck simulations, we varied onset (0.004, 0.04 and 0.4), strength (0.05, 0.1 and 0.5) and duration (0.08 and 0.4), and explored all possible combinations of those parameters. To compare different bottleneck scenarios, θ was scaled depending on the bottleneck parameters to keep SNP density constant for all simulations (on average ∼1850 SNPs per simulation). This was achieved by calculating a scaling factor (*f*) using the formula of Marth *et al.* (2004) and the approach described in DeGiorgio *et al.* (2014). The recombination rate was scaled to be 4*fN*_*e*_r to keep the mutation over recombination rate ratio comparable to the constant size simulations. The split time was also adjusted to *g*/*f*.

For the simulations with selective sweeps, we used 200 replicates for each parameter setting and sweep start time, and assumed *N*_*e*_ = 10,000. For calculation of the false positive rate, we conducted 4,000 neutral simulations under each bottleneck condition. For power calculations we generally assumed that the correct demographic model was known and used to identify critical values for the test, while for investigations of robustness we used the standard neutral model to estimate critical values. In all cases, the background site frequency spectrum was estimated using 1,000 neutral simulations. Note that in our analyses, the significance level is set so that 5% of all simulated 100kb regions are expected to contain at least one outlier, *i.e.* it is an experiment-wise significance level based on our simulated sequence length.

### Simulation of background selection

Background selection was simulated with the forward simulation software SFS_CODE (Hernandez 2008). To reduce the computational burden we simulated relatively small populations of *N*_e_ = 250 (Hernandez 2008). We used *n* = 15 and assumed constant population sizes with neutral and deleterious mutation rate of θ = 0.0025 per bp, *g*/(*4N*_*e*_) = 2, 4*N*_*e*_r = 0.15, and *L* = 100kb. We further assumed a selection coefficient of 2*N*_*e*_s = -50, reducing the neutral diversity by background selection by 40%. In the middle of the sequence (from 37.5kb to 62.5kb) we introduced a 100-fold reduction in recombination rate, which led to a local increase in the effect of background selection and an 80% reduction in SNP density (see Figure S1). This reduction in recombination rate mimics a selective sweep by locally reducing diversity through the effect of background selection (Figure S1). While the effect of background selection is more likely to act on a megabase scale (McVicker *et al.* 2009), we simulate strong background selection in a small segment of simulated sequence to keep the data sets small reducing the computational burden of the simulations. However, the difference in scale should not affect the generality of our conclusions.

To simulate selective sweeps in conjunction with background selection, a single positively selected mutation was introduced into the population 0.02 coalescence time units (2*N*_*e*_) in the past in the middle of the sequence, with a selection coefficient of 2*N*_*e*_s = 2000, or 0.1 coalescence time units in the past with a selection coefficient of 2*N*_*e*_*s* = 200. Whenever the mutation was lost from the simulation, the output was discarded and the simulation was repeated. For simulations without background selection, we set the deleterious mutation rate to zero. The composite likelihood ratio was calculated using a grid of 40 points for each simulated data set. The neutral simulations described above were used as background site frequency spectrum. For the HKA test we used non-overlapping windows of length 5kb.

### Analysis of human data

We used data from nine unrelated European individuals sequenced by Complete Genomics (Drmanac *et al.* 2010). Data and filtering steps were the same as in (DeGiorgio *et al.* 2014). We found that in low complexity regions around the centromeres and elsewhere in the genome, diversity drops to low levels while divergence to chimpanzee stays constant or even increases relative to other regions. CRG100 values, a measure of local alignability (Derrien *et al.* 2012), were downloaded from the UCSC Genome Browser at <u>http://genome.ucsc.edu/</u>. Low complexity regions are highly correlated with low values of CRG100 values, and increased levels of missing data. Therefore we only retained SNPs and fixed differences with a CRG value of 1 and full sample size. We also excluded windows with average CRG value of less than 0.9, in 100kb windows moving by 50kb.

We obtained recombination rates between pairs of sites from the sex-averaged pedigree-based human recombination map from deCODE Genetics (Kong *et al.* 2010). For the sweep scan, we calculated a composite likelihood ratio at grid points with 1kb spacing. We ran both standard SweepFinder, using only polymorphic sites (CLR1), and our new method using polymorphic sites, fixed differences relative to chimpanzees and the B-value map from McVicker et al (2009) (CLR2B). We assume an effective population size of humans and the human-chimpanzee ancestor population of 10,000 and 99,000, respectively, and a split time of 240,000 generations (McVicker *et al.* 2009). To look for overlaps with previous sweep scans, we use the supplementary table from (Akey 2009), compiling SFS based scans (Carlson *et al.* 2005; Kelley *et al.* 2006; Williamson *et al.* 2007), LD based scans (Wang *et al.* 2006; Voight *et al.* 2006; Kimura *et al.* 2007; Tang *et al.* 2007; Sabeti *et al.* 2007), and one *F*_*ST*_ based scan (Akey 2009).

## Results

### Including diversity as a sweep signal increases power and precision

We compare the power and accuracy of the CLR test when including only variable sites (CLR1), variable sites and fixed differences (CLR2), and all sites (CLR3), in the calculation of the composite likelihood ratio. CLR1 is the CLR that is calculated by current sweep detection software (Nielsen *et al.* 2005; Pavlidis *et al.* 2013). We start with a simple scenario of a constant population size with no background selection, and an advantageous mutation in the middle of the sequence, with selection strength of 2N_e_s = 200 and varying start times (see Methods).

The power drops quickly with the age of the selected mutation, and approaches zero for sweeps that start more than 0.5 coalescence time units (2*N*_*e*_ generations) in the past (Figure 3a). The root-mean-square error (RMSE) of the estimated location of the sweep also increases for older sweeps (Figure 3b). At an age of 0.5 coalescent time units, localization using the CLR1 statistic is not better than picking a site at random. In contrast, CLR2 and CLR3 still have power until 0.8 time units in the past. Furthermore, for sweeps that start 0.2 coalescence time units in the past, there is an almost 40% increase in power. In summary, both power and accuracy to localize the selected allele vastly increase when including fixed sites and there is little difference between including all sites (CLR3) and fixed differences (CLR2).

**Figure 3.**
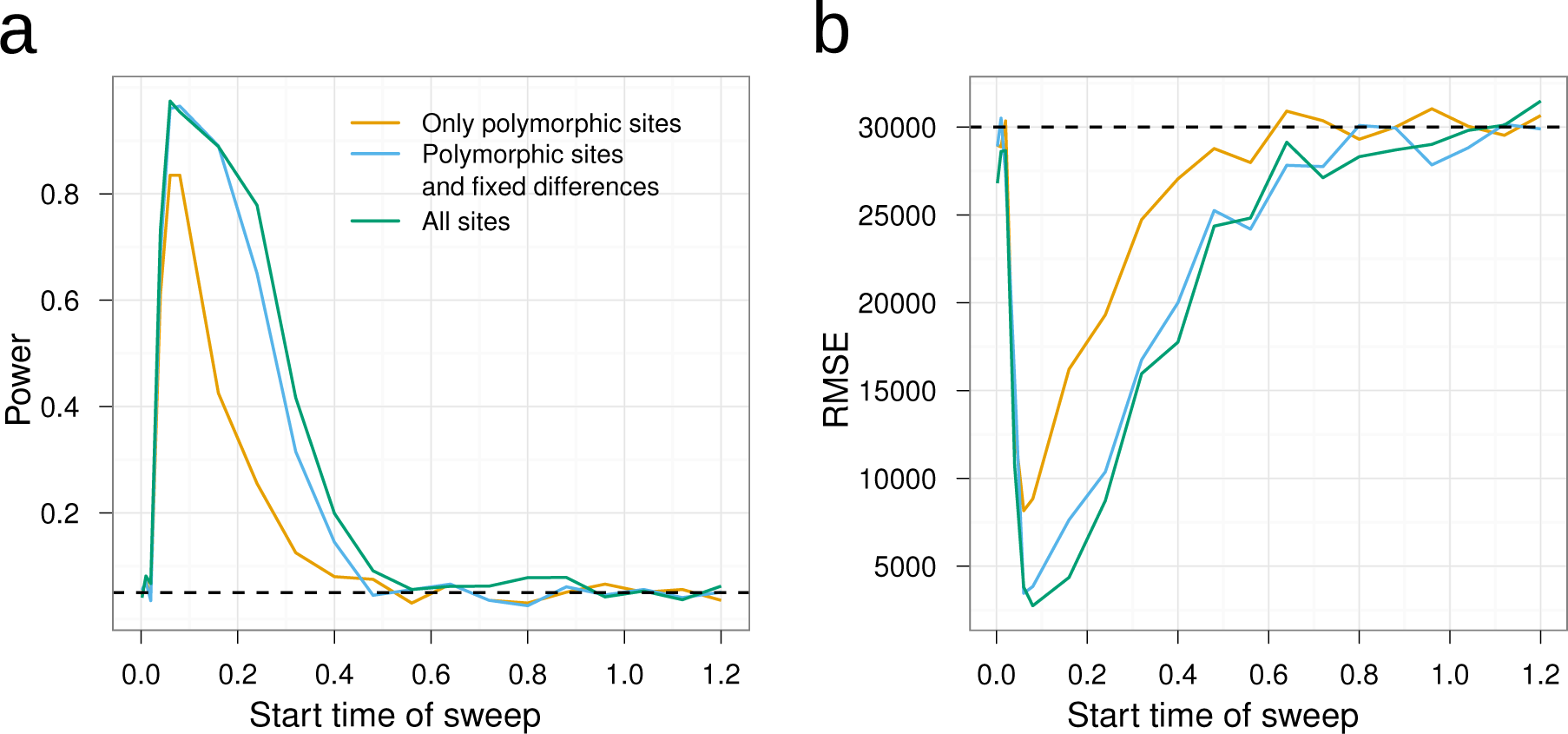
Power and accuracy comparison of the CLR tests. The power of the selection tests (a) and the root-mean-square error (RMSE) of the estimated location of the sweep (b) is shown as a function of the time since introduction of the beneficial mutation into the population in 2*N*_*e*_ generations (x-axis). The dashed line in (a) indicates the 5% significance level assumed in the power calculations, and in (b) it indicates the RMSE in case of random (uniform) localization of the sweep position. RMSE is calculated as the standard deviation of estimated minus true position in bp. Each 100kb simulated region is scored significant if it contains at least one significant outlier CLR at the 5% level.

### Including only fixed differences maintains robustness against mutation rate variation

We investigated the effect of varying mutation rates on the inference of sweeps. To this end we use two sets of simulations in 100 kb windows: one set with a population mutation rate of 0.005 and another set of simulations with reduced mutation rates relative to the first set. The likelihood ratio is then calculated using the first set of simulations as the background SFS when calculating the CLR for the second set (see Materials and Methods). The power is estimated by simulating a third set of simulations, with similarly varying mutation rates as in the second set, but with a beneficial mutations with selection coefficient 2*N*_*e*_*s* = 200 arising at 0.08 coalescence units in the past. The selected site is placed in the middle of the simulated region. In both cases, the null distribution of the test statistic is obtained using simulations with a constant high mutation rate of 0.005 and no selective sweeps.

If all sites are used for inference (CLR3), the power is close to 1 irrespective of the mutation rate. However, the false positive rate increases rapidly with the reduction in the mutation rate, so that at a 60% reduction already half the signals are false positives and at a reduction of 40%, almost all of the signals from the neutral simulations are false positives (Figure 4). This explains the apparently constant power. In contrast to CLR3, the power of both CLR1 and CLR2 reduces with the reduction in mutation rate (Figure 4). For CLR1, this reduction in power is due to the reduced SNP density. The power for CLR1 is only 80% to begin with and drops to 55% at a reduction in mutation rate by 50%. CLR2 is performing much better: the power to detect a sweep is still at 80% with a mutation rate reduction of 50%. The false positive rate for both CLR1 and CLR2 stays at or below the expected 5% level. In fact, the tests become extremely conservative when a mutation rate that is too high is used to obtain the null distribution of the composite likelihood ratio. This is because the distribution of the composite likelihood ratio is not invariant to the number of SNPs included in the analysis. Including many more SNPs for generating the null distribution (as a consequence of a higher mutation rate) than used in the analyses of the data, will result in a conservative test.

**Figure 4.**
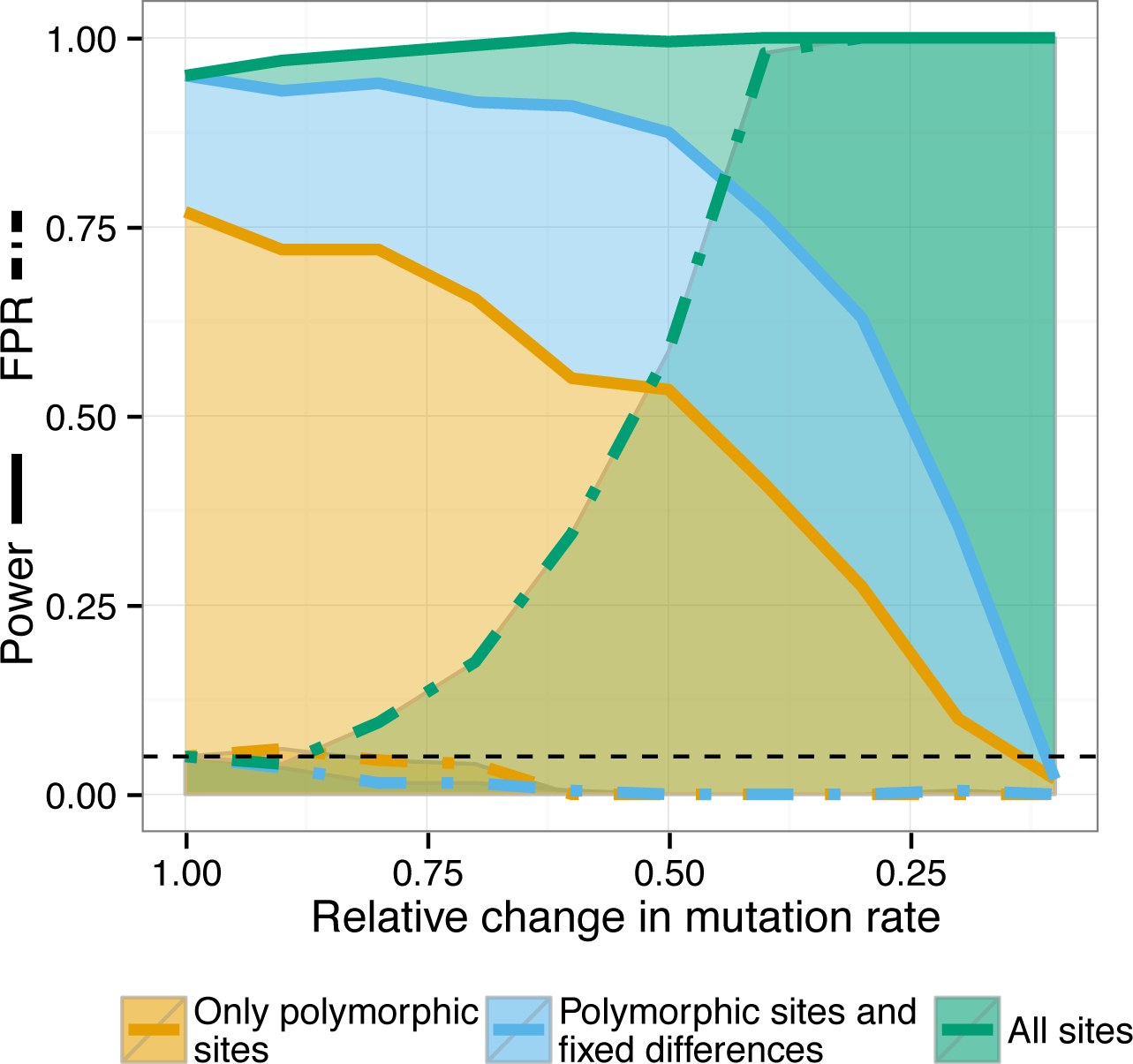
False positive rate (FPR) and power for reduced mutation rate. FPR and power, at a nominal significance level of 5%, as a function of the reduction in mutation rate of the sequence under investigation, relative to the mutation rate of the sequence that is used to calculate the background SFS. Both power and false positive rate are calculated by assuming a nominal significance level that is derived from simulations with no reduction in mutation rate (relative reduction = 1). Each 100kb simulated region is scored as significant if it contains at least one significant outlier CLR at the 5% level.

### Robustness to population bottlenecks

We simulated several bottleneck scenarios, varying onset, duration and strength of the bottleneck (Figure 5a), and calculated the false positive rates for the three sweep statistics (CLR1-3). Critical values for a 5% significance level were obtained from simulations with a constant size population. For each bottleneck scenario, we adjusted mutation rate, recombination rate and split time to the outgroup, so that the expected number of SNPs as well as divergence to the outgroup are comparable for all bottleneck models and for the constant size model (see Methods). This is equivalent to adjusting mutation rate and recombination rate in the simulations used to obtain critical values to match the observed data.

**Figure 5.**
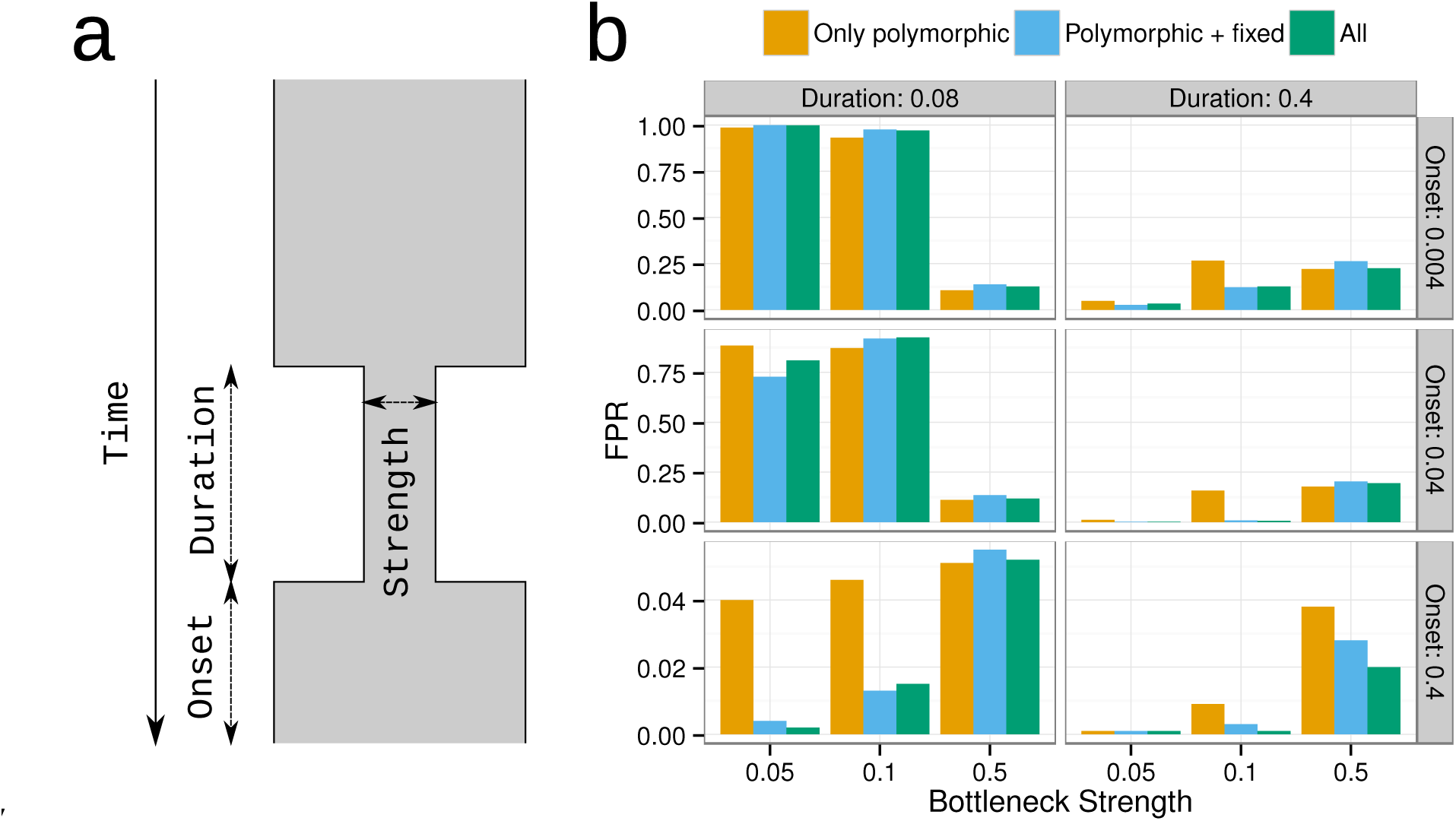
Robustness to population bottlenecks. a) Illustration of the bottleneck model used for the simulations, with varying onset time, duration and bottleneck strength leading to population size changes over time. ‘Strength’ is defined as *N*_*e*(*b*)_/*N*_*e*_ the effective population size during the bottleneck (*N*_*e*(*b*)_) divided by the effective population size before or after the bottleneck (*N*_*e*_), ‘duration’ is measured in number of generations divided by 2*N*_*e*_, and ‘onset’ is number of generations since the bottleneck started divided by 2*N*_*e*_. b) Proportion of false positives (probability of observing at least one wrongly inferred sweep) for bottleneck models if the null model for calculating statistical significance is based on a wrong constant size model with the same average number of SNPs and the same mutation to recombination ratio (see Methods for details). Each 100kb simulated region is scored significant if it contains at least one significant outlier CLR at the 5% level.

In a scenario with recent (onset = 0.004 or 0.04) and strong (strength = 0.05) or intermediate (strength = 0.1) bottlenecks, this generates a large proportion of false positives (>87%) if the population size is assumed to be constant (Figure 5b). The proportion of false positives is smaller if the bottleneck is old, as most lineages coalesce before or during the bottleneck. This is true for all three CLR statistics. However, by including invariant sites or fixed differences into the CLR framework, we increase robustness to bottlenecks whenever the chance of surviving the bottleneck is relatively small, *e.g.* when the bottleneck is strong (5%), or intermediate (10%) and has a long duration (0.2).

As a specific example, European humans are assumed to have experienced a bottleneck during colonization of Europe. Estimated bottleneck parameters (Lohmueller *et al.* 2011b) indicate a relatively recent, short, but strong bottleneck (onset = 0.014, duration = 0.005, strength = 0.05). Simulating data under this scenario results in a proportion of false positives of 0.21 for CRL1 and CLR2 and 0.24 for CLR3, suggesting that constant population size is not a suitable demographic model for calculating significance thresholds for any of the three CLR tests.

### False positives due to background selection are prevented by including a *B*-value map

A strong reduction in diversity relative to divergence in regions of the genome can be caused not only by selective sweeps, but also by the effects of deleterious mutations on linked neutral variation, *i.e.* background selection. We adapted SweepFinder to enable the inclusion of genome-wide estimates of this effect, the B-value map, to account for this type of variation. To evaluate the method, we simulated a genomic region with increased background selection, *i.e.* a local reduction in diversity due to background selection (Figure S1, Figure 6 and Methods). The background SFS used to calculate the CLR statistics was based on, otherwise identical, neutral simulations. To evaluate power in the presence of background selection, we simulated data with both background selection and a recently completed selective sweep located in the middle of the sequence (Figure 6). The nominal false positive rate, which is used to determine the nominal significance level, was estimated from neutral simulations without background selection.

**Figure 6.**
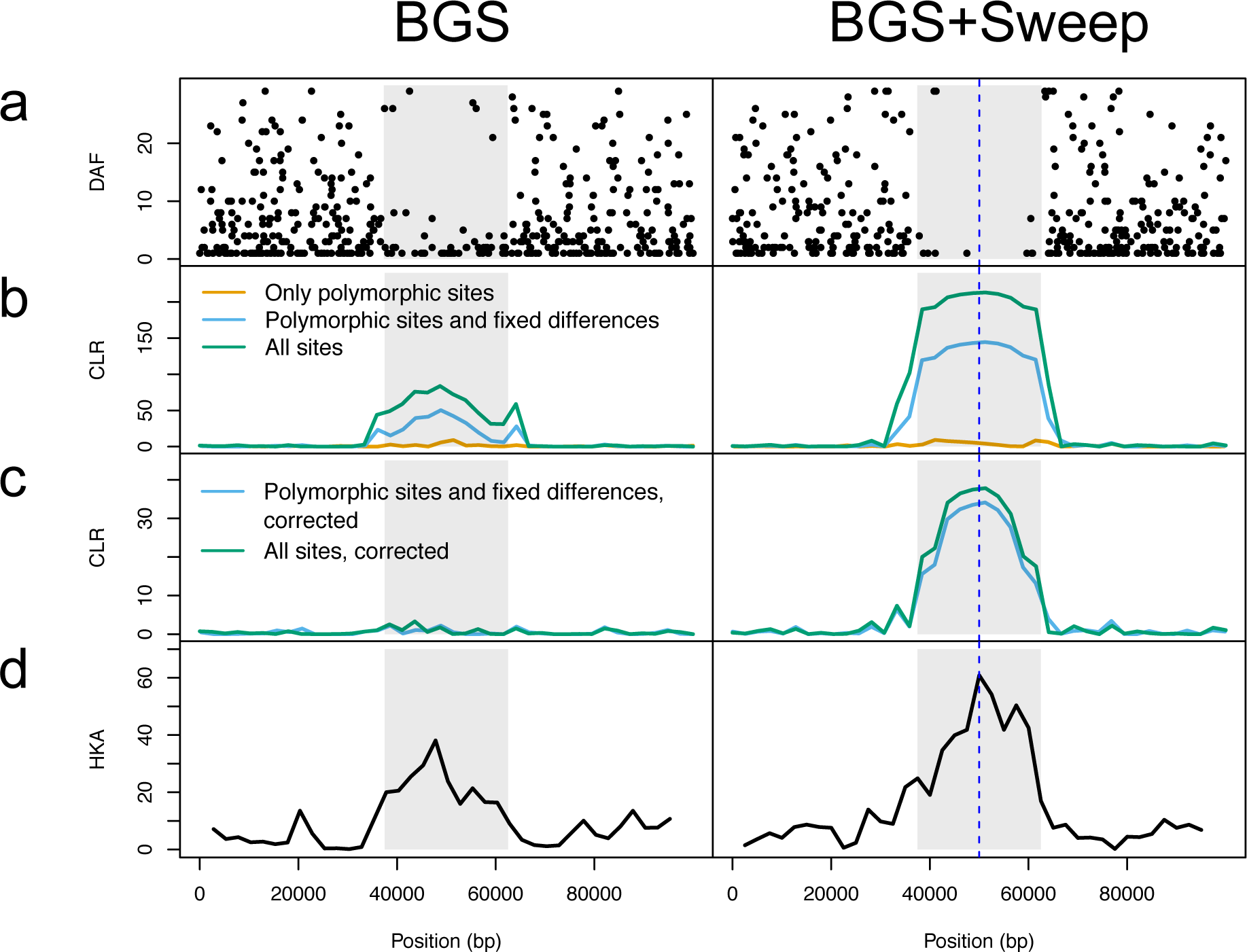
Two examples of simulation results from forward simulations. Plotted are a) the derived allele frequency (DAF) of each SNP across the 100kb sequence, b) CLR without correction for background selection, c) CLR corrected for background selection, d) HKA test statistic (signed chi-square statistic in non-overlapping windows). There is a uniform deleterious mutation rate across the 100kb sequence. A 100-fold reduction in recombination rate in the middle part of the sequence (grey box) generates a larger background selection effect in that part compared to the surrounding sequence (see also Figure S1). Left: Only background selection (BGS). Right: Background selection together with a recently fixed selective sweep in the middle of the sequence (BGS+Sweep).

The HKA test and the uncorrected CLR2 and CLR3 cannot distinguish background selection from selective sweeps, as is evident from the nearly 100% false positives under our strong background selection scenario (Figure 7a). If only polymorphic sites (CLR1) are used, the test does not suffer from an elevated level of false positives, indicating that CLR2 and CLR3 mainly pick up on the diversity reduction. However, if the diversity reduction due to background selection is factored in using a B-value map, the statistics return to the desired behavior in that the FPR corresponds to the nominal significance level, while maintaining increased power as compared to CLR1.

**Figure 7.**
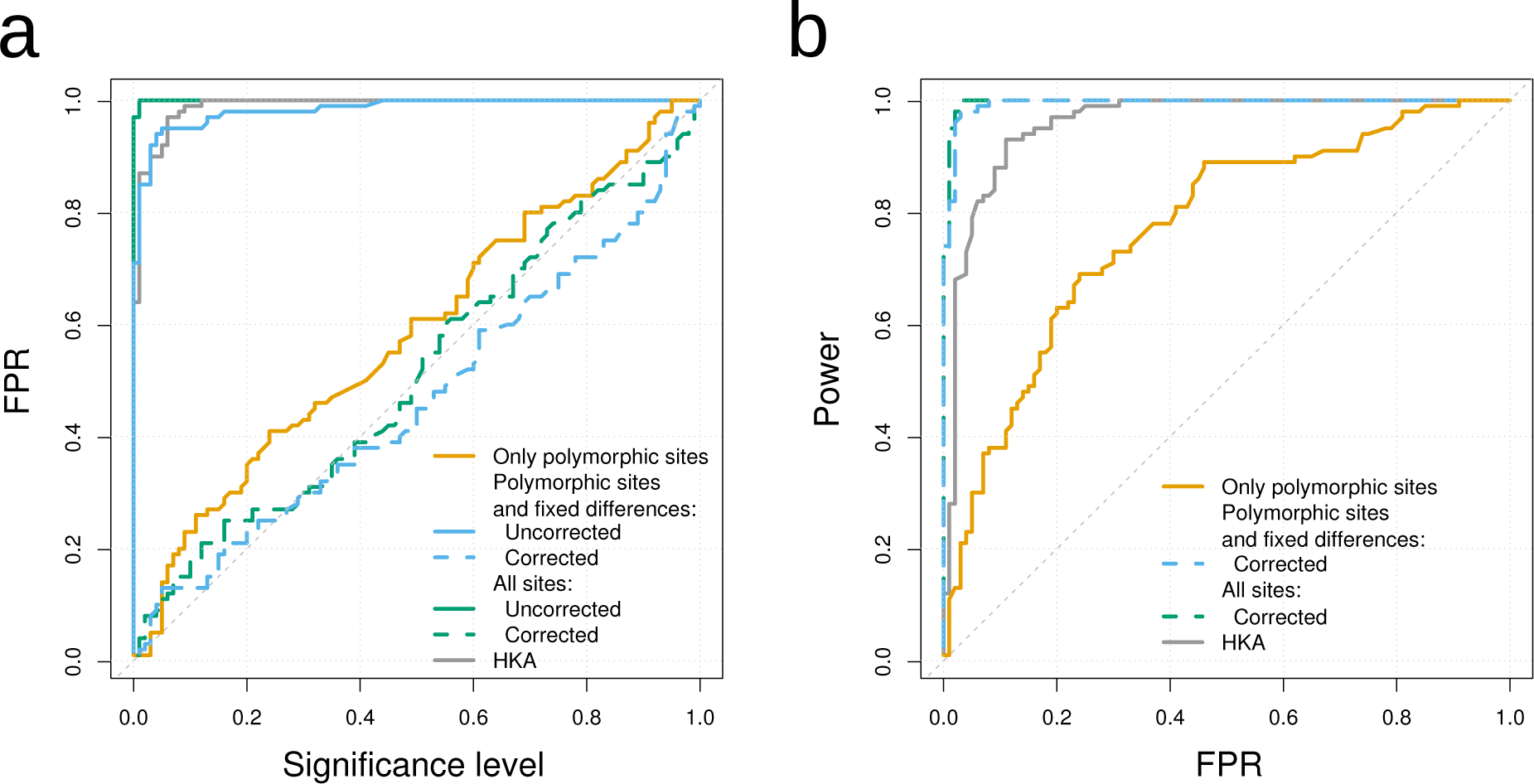
FPR and power under background selection. a) The observed proportion of false positives in case of simulations with background selection plotted against the nominal false positive rate (significance level). The nominal false positive rate is estimated from neutral simulations without background selection. b) The power to detect a recently fixed selective sweep with 2*N*_*e*_*s* = 2000 as a function of the proportion of false positives (see supplemental Figure S5b for results with 2*N*_*e*_*s*=200).

### Analysis of a human genetic variation dataset

We screened the data from nine unrelated European individuals sequenced by Complete Genomics (Drmanac *et al.* 2010) for selective sweeps to prove the utility of our improvements to SweepFinder. We compare the composite likelihood ratio across the whole genome, calculated using only polymorphic sites (CLR1), with our new approach by including fixed differences to chimpanzees into the calculation (CLR2). To account for varying diversity across the genome due to background selection we also incorporate the B-value map from McVicker et al. (2009) into the calculation of CLR2, henceforth referred to as CLR2B.

Due to the complex human demography and the added complication of background selection, we do not calculate critical values, but report the 0.2% most extreme regions in Table 1. This approach has previously been used in other selection scans (e.g., Voight *et al.* 2006) under the argument that it is an outlier approach, although we notice that no formal testing has been done here or in Voight *et al.* (2006) to determine the degree to which the most extreme values indeed are outlying with respect to some parametric distribution.

**Table 1.**
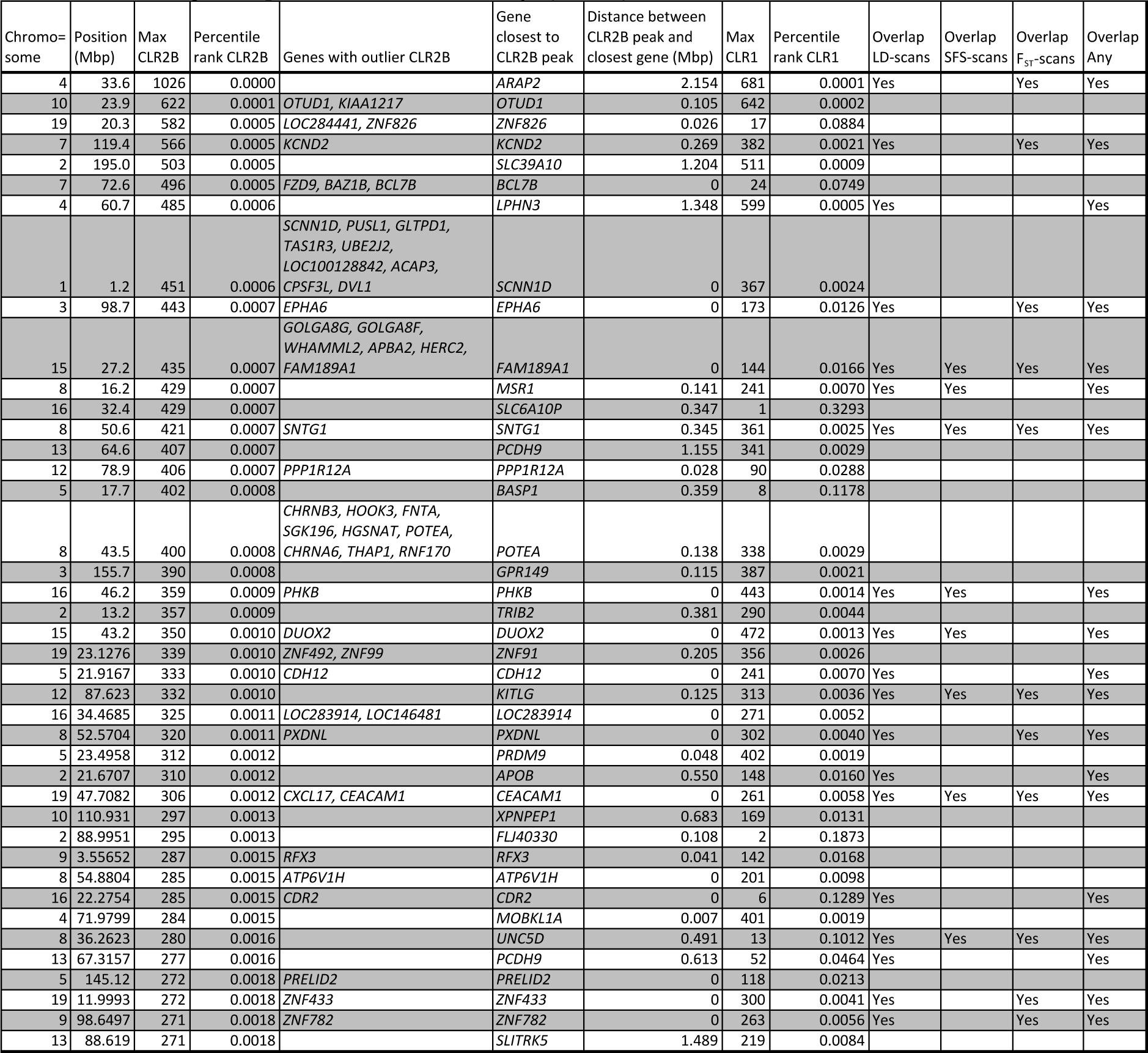
A list of sweep regions, using an outlier approach. Only regions with CLR2B values larger than the genome-wide 99.8% quantile are shown. Consecutive outlier CLR2B values are merged to a single sweep region. The overlap with previous scans is tabulated using compiled data from Akey (2009).

The strongest sweep signal is on chromosome 4, 33.6 Mbp, a region without any annotated genes. The closest gene, *ARAP2*, is 2.15 Mbp downstream from the CLR2 peak. This sweep region has a B-value close to one, and a strong reduction in diversity relative to divergence. The peak in CLR1 shows that this region is characterized by a sweep-like site frequency spectrum. This region was also listed as a candidate region in LD-based (Wang *et al.* 2006; Voight *et al.* 2006; Kimura *et al.* 2007; Sabeti *et al.* 2007) and SFS- based sweep scans (Carlson *et al.* 2005; Kelley *et al.* 2006; Williamson *et al.* 2007).

The gene with the strongest CLR2B signal is *KIAA1217*, which was suggested to affect lumbar disc herniation susceptibility (Karasugi *et al.* 2009). The gene is also an outlier for haplotype-based sweep statistic for detecting incomplete soft or hard sweeps, in an African population (Ferrer-Admetlla *et al.* 2014). This may suggest that the variant is fixed, or at very high frequency in Europe, but still polymorphic in Africa. Another gene in one of the outlier regions, *HERC2*, is known to modulate iris color and blonde hair (Wilde *et al.* 2014). This candidate has previously been identified in a screen for population-specific sweeps using XP-CLR (Chen et al. 2010). Analyses of ancient DNA suggests that strong selection has been operating on *HERC2* in western Eurasia during the past 5000 years (Wilde *et al.* 2014).

About half of our outlier regions in Table 1 overlap with at least one candidate region of previous sweep scans in humans (Akey 2009), and most of them are also outlier regarding CLR1. However, there are some notable exceptions: One example is the sweep region on chromosome 7, at 72.6 Mbp, with the genes *BCL7B*, *FZD9* and *BAZ1B*. This region has a small CLR2B percentile rank of 0.0005, but a much larger CLR1 percentile rank (0.071), and is not listed in Akey (2009).

In conclusion, we show that CLR2B shows enrichment for previously detected candidates, but also identifies novel sweep signals. These previously undetected sweeps are likely to be enriched for sweeps that started between 0.2 and 0.8 *N*_e_ generations ago and thus escaped detection with LD, *F*_*ST*_ or SFS based methods.

## Discussion

We evaluated the performance of a composite likelihood ratio test for detecting selection sweeps (Nielsen *et al.* 2005) when including fixed differences in the likelihood ratio in addition to SFS information, using extensive simulations. We show that there can be a marked increase in power as well as a reduction in false positive rate for a number of different scenarios in several different models of mutation rate variation, population bottlenecks and background selection. We also show that estimates of the strength of background selection can be included into the framework to prevent false positives in regions with strong, long-term background selection. By applying the method to human genetic data, we detect novel regions that are not identified as candidate regions with the standard SweepFinder approach.

### Using invariant sites increases power and robustness

Given that both diversity and divergence change proportionally with mutation rate, we integrate variation in mutation rates by including a measure of divergence to an outgroup species. More specifically, we include sites that are not polymorphic within the species under investigation, but differs from an outgroup sequence, i.e. inferred fixed differences. If the SweepFinder CLR is calculated including all sites, ignoring outgroup information, variation in mutation rates can create false positives (Figure 4). However, if only fixed differences are added to the SFS, the power, but not the false positive rate, increases.

Furthermore, including invariant sites can increase robustness to certain bottleneck scenarios if the bottleneck is of intermediate to high strength, but not too recent (Boitard *et al.* 2009; Pavlidis *et al.* 2010). However, like many other methods for detecting selective sweeps (Barton 1998; Jensen *et al.* 2005; Voight *et al.* 2006; Boitard *et al.* 2009; Pavlidis *et al.* 2010; Crisci *et al.* 2013) the CLR test can suffer from a disturbingly high false positive rate in the presence of recent bottlenecks in population size. The use of an empirically derived demographic background SFS does not eliminate the sensitivity to demographic assumptions, because the CLR does not model the correlation in coalescence times along the sequence correctly irrespective of the demographic model. A bottleneck will force many lineages to coalesce in a short amount of time. If the duration of the bottleneck is such that at least some lineages escape the bottleneck in most regions, the few regions in which all lineages coalesce during the bottleneck may very much resemble regions that have been affected by a selective sweep. Realistic demographic models should be used if assigning p-values to individual sweeps.

### Background selection as a null model for sweep detection

What is often neglected in previous discussions of diversity-based sweep detection methods is variation in diversity across the genome that is not caused by variation in mutation rate (or conservation level), but by variation in background selection, *i.e.* by the effect of deleterious mutations on linked neutral variation (Charlesworth *et al.* 1993; Hudson & Kaplan 1995; Charlesworth 2012; Cutter & Payseur 2013). A locally increased level of background selection will lead to a reduction in diversity similar to that expected after a selective sweep.

As datasets and methods for estimating the effect of background selection for each position in the genome are becoming available (McVicker *et al.* 2009), the objective of developing methods for detecting positive selection that can take background selection into account is becoming tenable. We present the first such method by including a map of predicted B-values in the calculation of the CLR. McVicker *et al.* (2009) provide such a B-value map for humans by defining functional elements based on mammalian sequence conservation, and fitting parameters to phylogenetic data. Therefore, reductions in neutral diversity in regions of the human data do not influence the local estimation of *B*. Our approach considers a local reduction in diversity as evidence for a selective sweep only if it is not also predicted by a local drop in B-values, *i.e.* background selection is our evolutionary null model (Cutter & Payseur 2013). We simulated background selection levels typical for humans (McVicker *et al.* 2009), and by accounting for background selection, we could effectively prevent false positives without loosing power. If one does not account for background selection, the proportion of false positives is large and similar to that of a HKA test (Figure 7a).

### Application to human data

Finally, by applying our method to human genetic variation data, we show that the new method detects novel regions that were not identified as candidates using the standard SweepFinder approach. Based on our simulations, we would expect those regions to be enriched for old selective sweeps that started between 0.2 and 0.8 *N*_*e*_ generations ago, a time range where the power of other SFS-based, *F*_ST_ and LD based methods is low (Sabeti *et al.* 2006). Interestingly, the strongest signal we find, which has been missed by most previous scans, is near *KIAA1217*, a gene affecting lumbar disc herniation susceptibility. We speculate that the selection in this region may possibly be related to changes in human muscular-skeletal function subsequent to the evolution of erect bipedal walk. Increased risk of lumbar disc herniation is a likely consequence of bipedal walk. We may still be evolving to optimize muscular–skeleton functions after this recent, radical change in skeletal structure and function.

## Acknowledgments

We gratefully acknowledge Melissa Hubisz for her help with the original SweepFinder software. This work was supported by the Austrian Science Fund (Vienna Graduate School of Population Genetics, FWF W1225) to C.D.H.

## Data Accessibility

We did not generate any new data set for this study.

## Author Contributions

CDH, IH and RN conceived the study question and the experimental design. CDH implemented the simulations. CDH and MD adapted the software SweepFinder to account for background selection. CDH and MD analyzed the simulations and the human genetic variation data. CDH, MD, IH and RN wrote the paper.

## References

Akey JM (2009) Constructing genomic maps of positive selection in humans: Where do we go from here? Genome Research, 19, 711–722.

Akey JM, Zhang G, Zhang K, Jin L, Shriver MD (2002) Interrogating a high-density SNP map for signatures of natural selection. Genome Research, 12, 1805–1814.

Barton NH (1998) The effect of hitch-hiking on neutral genealogies. Genetical Research,72, 123–133.

Boitard S, Schlotterer C, Futschik A (2009) Detecting selective sweeps: a new approach based on hidden Markov models. Genetics, 181, 1567.

Carlson CS, Thomas DJ, Eberle MA et al. (2005) Genomic regions exhibiting positive selection identified from dense genotype data. Genome Research, 15, 1553–1565.

Charlesworth B (2012) The Effects of Deleterious Mutations on Evolution at Linked Sites. Genetics, 190, 5–22.

Charlesworth D, Charlesworth B, Morgan MT (1995) The pattern of neutral molecular variation under the background selection model. Genetics, 141, 1619–1632.

Charlesworth B, Morgan MT, Charlesworth D (1993) The effect of deleterious mutations on neutral molecular variation. Genetics, 134, 1289–1303.

Chávez-Galarza J, Henriques D, Johnston JS et al. (2013) Signatures of selection in the Iberian honey bee (Apis mellifera iberiensis) revealed by a genome scan analysis of single nucleotide polymorphisms. Molecular Ecology, 22, 5890–5907.

Chen H, Patterson N, Reich D (2010) Population differentiation as a test for selective sweeps. Genome Research, 1–10.

Comeron JM (2014) Background Selection as Baseline for Nucleotide Variation across the Drosophila Genome. PLoS Genet, 10, e1004434.

Crisci JL, Poh Y-P, Mahajan S, Jensen JD (2013) The impact of equilibrium assumptions on tests of selection. Evolutionary and Population Genetics, 4, 235.

Cutter AD, Payseur BA (2013) Genomic signatures of selection at linked sites: unifying the disparity among species. Nature reviews. Genetics, 14, 262–274.

DeGiorgio M, Lohmueller KE, Nielsen R (2014) A Model-Based Approach for Identifying Signatures of Ancient Balancing Selection in Genetic Data. PLoS Genet, 10, e1004561.

Derrien T, Estellé J, Marco Sola S et al. (2012) Fast Computation and Applications of Genome Mappability. PLoS ONE, 7, e30377.

Drmanac R, Sparks AB, Callow MJ et al. (2010) Human Genome Sequencing Using Unchained Base Reads on Self-Assembling DNA Nanoarrays. Science, 327, 78–81.

Durrett R, Schweinsberg J (2004) Approximating selective sweeps. Theoretical population biology, 66, 129–138.

Ewing G, Hermisson J (2010) MSMS: A Coalescent Simulation Program Including Recombination, Demographic Structure, and Selection at a Single Locus. Bioinformatics (Oxford, England), 26, 2064–2065.

Ferrer-Admetlla A, Liang M, Korneliussen T, Nielsen R (2014) On Detecting Incomplete Soft or Hard Selective Sweeps Using Haplotype Structure. Molecular Biology and Evolution, 31, 1275–1291.

Fu YX, Li WH (1993) Statistical tests of neutrality of mutations. Genetics, 133, 693–709.

Garud NR, Messer PW, Buzbas EO, Petrov DA (2015) Recent Selective Sweeps in North American Drosophila melanogaster Show Signatures of Soft Sweeps. PLoS Genet,11, e1005004.

Hernandez RD (2008) A flexible forward simulator for populations subject to selection and demography. Bioinformatics, 24, 2786–2787.

Huber CD, Nordborg M, Hermisson J, Hellmann I (2014) Keeping It Local: Evidence for Positive Selection in Swedish Arabidopsis thaliana. Molecular Biology and Evolution, 31, 3026–3039.

Hudson RR, Kaplan NL (1994) Gene Trees with Background Selection. In: Non-Neutral Evolution (ed Golding B), pp. 140–153. Springer US.

Hudson RR, Kaplan NL (1995) Deleterious background selection with recombination. Genetics, 141, 1605–1617.

Hudson RR, Kreitman M, Aguadé M (1987) A Test of Neutral Molecular Evolution Based on Nucleotide Data. Genetics, 116, 153–159.

Jensen JD, Kim Y, DuMont VB, Aquadro CF, Bustamante CD (2005) Distinguishing Between Selective Sweeps and Demography Using DNA Polymorphism Data. Genetics, 170, 1401–1410.

Jensen JD, Thornton KR, Bustamante CD, Aquadro CF (2007) On the Utility of Linkage Disequilibrium as a Statistic for Identifying Targets of Positive Selection in Nonequilibrium Populations. Genetics, 176, 2371–2379.

Karasugi T, Semba K, Hirose Y et al. (2009) Association of the tag SNPs in the human SKT gene (KIAA1217) with lumbar disc herniation. Journal of Bone and Mineral Research: The Official Journal of the American Society for Bone and Mineral Research, 24, 1537–1543.

Kelley JL, Madeoy J, Calhoun JC, Swanson W, Akey JM (2006) Genomic signatures of positive selection in humans and the limits of outlier approaches. Genome Research, 16, 980–989.

Kim Y, Nielsen R (2004) Linkage Disequilibrium as a Signature of Selective Sweeps. Genetics, 167, 1513–1524.

Kim Y, Stephan W (2002) Detecting a local signature of genetic hitchhiking along a recombining chromosome. Genetics, 160, 765–777.

Kimura R, Fujimoto A, Tokunaga K, Ohashi J (2007) A Practical Genome Scan for Population-Specific Strong Selective Sweeps That Have Reached Fixation. PLoS ONE, 2, e286.

Kong A, Thorleifsson G, Gudbjartsson DF et al. (2010) Fine-scale recombination rate differences between sexes, populations and individuals. Nature, 467, 1099–1103.

Lewontin RC, Krakauer J (1973) Distribution of gene frequency as a test of the theory of the selective neutrality of polymorphisms. Genetics, 74, 175–95.

Li H (2011) A New Test for Detecting Recent Positive Selection that is Free from the Confounding Impacts of Demography. Molecular Biology and Evolution, 28, 365–375.

Lohmueller KE, Albrechtsen A, Li Y et al. (2011a) Natural Selection Affects Multiple Aspects of Genetic Variation at Putatively Neutral Sites across the Human Genome. PLoS Genet, 7, e1002326.

Lohmueller KE, Bustamante CD, Clark AG (2011b) Detecting Directional Selection in the Presence of Recent Admixture in African-Americans. Genetics, 187, 823–835.

Long Q, Rabanal FA, Meng D et al. (2013) Massive genomic variation and strong selection in Arabidopsis thaliana lines from Sweden. Nature Genetics, 45, 884–890.

Marth GT, Czabarka E, Murvai J, Sherry ST (2004) The allele frequency spectrum in genome-wide human variation data reveals signals of differential demographic history in three large world populations. Genetics, 166, 351–372.

McVicker G, Gordon D, Davis C, Green P (2009) Widespread Genomic Signatures of Natural Selection in Hominid Evolution. PLoS Genet, 5, e1000471.

Messer PW, Petrov DA (2013) Frequent adaptation and the McDonald-Kreitman test. Proceedings of the National Academy of Sciences of the United States of America, 110, 8615–8620.

Nicolaisen LE, Desai MM (2013) Distortions in Genealogies due to Purifying Selection and Recombination. Genetics, 195, 221–230.

Nielsen R, Williamson S, Kim Y et al. (2005) Genomic scans for selective sweeps using SNP data. Genome Research, 1566–1575.

Nordborg M, Charlesworth B, Charlesworth D (1996) The effect of recombination on background selection. Genetics Research, 67, 159–174.

Pavlidis P, Hutter S, Stephan W (2008) A population genomic approach to map recent positive selection in model species. Molecular Ecology, 17, 3585–3598.

Pavlidis P, Jensen JD, Stephan W (2010) Searching for footprints of positive selection in whole-genome SNP data from nonequilibrium populations. Genetics, 185, 907–22.

Pavlidis P, Živković D, Stamatakis A, Alachiotis N (2013) SweeD: Likelihood-Based Detection of Selective Sweeps in Thousands of Genomes. Molecular Biology and Evolution, 30, 2224–2234.

Qanbari S, Strom TM, Haberer G et al. (2012) A High Resolution Genome-Wide Scan for Significant Selective Sweeps: An Application to Pooled Sequence Data in Laying Chickens. PLoS ONE, 7, e49525.

Ramey HR, Decker JE, McKay SD et al. (2013) Detection of selective sweeps in cattle using genome-wide SNP data. BMC Genomics, 14, 382.

Sabeti PC, Reich DE, Higgins JM et al. (2002) Detecting recent positive selection in the human genome from haplotype structure. Nature, 419, 832–837.

Sabeti PC, Schaffner SF, Fry B et al. (2006) Positive Natural Selection in the Human Lineage. Science, 312, 1614–1620.

Sabeti PC, Varilly P, Fry B et al. (2007) Genome-wide detection and characterization of positive selection in human populations. Nature, 449, 913–8.

Tang K, Thornton KR, Stoneking M (2007) A New Approach for Using Genome Scans to Detect Recent Positive Selection in the Human Genome. PLoS Biol, 5, e171.

Voight BF, Kudaravalli S, Wen X, Pritchard JK (2006) A Map of Recent Positive Selection in the Human Genome. PLoS Biol, 4, e72.

Wang ET, Kodama G, Baldi P, Moyzis RK (2006) Global landscape of recent inferred Darwinian selection for Homo sapiens. Proceedings of the National Academy of Sciences of the United States of America, 103, 135–140.

Wilde S, Timpson A, Kirsanow K et al. (2014) Direct evidence for positive selection of skin, hair, and eye pigmentation in Europeans during the last 5,000 y. Proceedings of the National Academy of Sciences, 201316513.

Williamson SH, Hubisz MJ, Clark AG et al. (2007) Localizing recent adaptive evolution in the human genome. PLoS genetics, 3, e90.

Williford A, Comeron JM (2010) Local effects of limited recombination: historical perspective and consequences for population estimates of adaptive evolution. The Journal of Heredity, 101 Suppl 1, S127–134.

Xia Q, Guo Y, Zhang Z et al. (2009) Complete resequencing of 40 genomes reveals domestication events and genes in silkworm (Bombyx). Science (New York, N.Y.), 326, 433–436.

Zeng K, Charlesworth B (2011) The joint effects of background selection and genetic recombination on local gene genealogies. Genetics, 189, 251–266.

